# Assessing accelerometry as an alternative to electromyography for mouse sleep scoring

**DOI:** 10.64898/2026.09.25.754505

**Authors:** Dawson Green, Justine Fortin-Houde, Elsa Lalot, Nicolas Lemmetti, Tanya Leduc, Alice Gravel-Chouinard, Guillaume Ducharme, Valérie Mongrain, Bénédicte Amilhon

## Abstract

For sleep scoring in rodents, the use of electromyography (EMG) is considered the gold standard to assess muscle tone; however, EMG electrodes require invasive surgical implantation, are sensitive to muscular placement, can have extensive noise due to non-muscular signal contamination, and are at risk of failure due to mechanical stress. The use of an accelerometer (Acc) integrated on the recording headstage offers an alternative for movement detection. However, the degree of agreement between these two approaches has never been systematically evaluated. To understand if these tools allow similar scoring outcomes, seven trained scorers manually classified mouse sleep recordings using hippocampal local field potential (LFP) together with EMG or Acc. When comparing scoring between EMG and Acc, overall epoch-by-epoch accuracy exceeded 90% with a kappa score above 0.8, indicating excellent agreement. However, Acc was found to under classify Wake by 4.92% while overclassifying NREM sleep by 3.29%, possibly reflecting a decreased sensitivity of Acc to subtle changes in muscle tone used in distinguishing Wake from NREM sleep. Comparing EMG and Acc signal for agreed epochs vs disagreed epochs revealed a weaker correlation between the two signals for disagreed epochs, especially for Wake and NREM, suggesting that disagreements reflected genuine signal differences rather than scorer subjectivity. These findings support LFP/Acc as a valid approach to sleep scoring, however, it is noted that Acc based scoring slightly underestimated Wake epochs, favoring NREM sleep, relative to traditional EMG approach.

## INTRODUCTION

Sleep is an important and foundational physiological process essential for maintaining cognitive functioning, metabolic balance, emotional stability, and overall health, with the necessity of sleep extending across virtually every organ system^1,2^. Disrupted and inadequate sleep has been linked with a broad range of human pathologies including cardiovascular disease, metabolic dysfunction, and neuropsychiatric disorders^3,4^. Thus, sleep has become an important focus of both basic and translational neuroscience research.

Rodent models, particularly mice, have played a crucial role in furthering our understanding of sleep. Sleep has largely been assessed using a combination of electroencephalography (EEG) and electromyography (EMG) to respectively capture changes in cortical activity and muscle tone across vigilance states^5-8^. Local field potentials (LFP) that capture intracranial electrical activity can be used just as EEG to assess sleep and wake states in combination with EMG^8^. Together, LFP and EMG allow classification into three different generally accepted states; wakefulness, non-rapid eye movement (NREM) sleep, and rapid eye movement (REM) sleep, all reflecting distinct waveform signatures and muscular activity patterns^8-10^. This process of sleep scoring has become increasingly automated through deep learning-based algorithms and automatic scoring programs, further enhancing the efficiency, throughput, and reproducibility of sleep analysis^11–20^. However, manual scoring by trained experts remains the gold standard in many research settings, particularly when nuanced state identification is required or when validating automated approaches^21^. Despite continued refinement of EEG or LFP combined with EMG-based sleep scoring, this approach still carries inherent limitations. Surgical implantation of EMG electrodes is invasive, technically demanding, requires substantial recovery time, is sensitive to muscular placement, can have extensive noise due to non-muscular signal contamination, and are at risk of failure due to mechanical stress^22^. These challenges can be particularly significant in affecting data quality and therefore scoring outcomes.

As a result, accelerometers (Acc) have gained increasing interest as a lower burden alternative^23-26^. Acc, which detects gross body movement and is integrated directly into a recording head stage, offers a substitute for muscle activity, without requiring a separate surgical implantation site. Acc sensors measure acceleration, converting animal movement into a voltage signal proportional to acceleration. Because movements produce distinctive, time-varying acceleration signatures across x, y, and z axes, this signal could be used as a substitute for EMG in distinguishing muscular activity. We therefore can predict Acc and EMG signal to be highly correlated, which offers the opportunity for Acc utilization in sleep scoring. However, this raises important questions regarding the accuracy, reliability, and potential systematic biases in the classification of vigilance states through Acc. Here, we used intracranial LFP recordings where both EMG and Acc were simultaneously recorded in mice and compared the manual sleep scoring from trained scorers between modalities to present the first direct comparison of LFP/EMG and LFP/Acc combinations in the identification of mouse wake/sleep stages.

## METHODS

### Animals

All procedures involving animals were approved by the Comité Institutionel de Bonnes Pratiques Animales en Recherche (CIBPAR) of the CHU Sainte-Justine Azrieli Research Center, in accordance with the standards of the Canadian Council on Animal Care. The animals and recordings presented in this paper were originally part of another study (unpublished results). In short, heterozygous SERT-CRE mice (n = 6; MMRRC#031028-UCD) were injected in the median raphe ([AP] -4.48, [ML] +/-0.04 at 10° angle, [DV] -4.36) with one of the following viral constructs: AAVdj-EF1a-DIO hChR2E123T/T159C eYFP or AAVdj-EF1a-DIO eYFP. Both male (n = 5) and female (n = 1) were used. After surgery, mice were housed individually on a 12:12-hour light/dark cycle, 24°C, 50% humidity and with ad libitum access to food and water.

### Surgical Implantation

Implants consisted of a single 100 µm tungsten wire electrode (A-M Systems), a stainless-steel wire electrode for EMG recording (A-M Systems), and a skull screw placed over the cerebellum (Antrin Miniature Specialties) serving as both ground and reference. Mice were anesthetized with isoflurane (5% induction, 1–2% maintenance) and placed in a stereotaxic frame. After exposure and cleaning of the skull, small craniotomies were drilled, and the electrode was implanted in the right dorsal hippocampal CA1 ([AP] -2.25 mm, [ML] +2.25 mm, [DV] -1.80 mm relative to bregma). The EMG electrode was inserted into the dorsal neck (trapezius) muscles. The implant was secured with C&B Metabond (Parkell) and dental acrylic (Patterson Dental). Mice received carprofen (20 mg/kg) for analgesia and saline for rehydration. Mice were allowed at least 2 weeks of recovery before recordings.

### Recordings

After recovery, mice were habituated to handling and connection cabling with a sham headstage for three to five days. LFP recordings were performed in the home cage during the light period (ZT3.5 to ZT10) and between five and fourteen weeks post-surgery. Mouse implants were connected to a 32-channel headstage (Open Ephys SPI Headstage 32ch, SKU: OEPS-6573 [Accelerometer]) linked to the Open Ephys acquisition system. LFP and EMG signals were digitised at 5 kHz. The headstage-integrated accelerometer provided head movement data by measuring acceleration across x, y, and z axes. LFP, Acc and EMG recordings were recorded simultaneously, providing 40.21 hours of simultaneous LFP, Acc and EMG data. This yielded 80.5 total hours of data to sleep score (n = 6 mice), with an average of 6.7 hours per mouse.

### Sleep Scoring

Vigilance states were manually scored by seven trained scorers using both LFP/EMG and LFP/Acc recordings in a custom MATLAB-based interface. Recordings were segmented into 4-second epochs and classified into Wake, NREM sleep, and REM sleep. Classification followed standard rodent sleep scoring criteria^5-10,27-29^. In short, Wake was characterized by low-amplitude, high frequency LFP activity together with high and variable EMG tone or Acc signal amplitude; NREM sleep by high amplitude, low frequency LFP activity with low and stable EMG tone or Acc signal amplitude; and REM sleep by low amplitude, theta-enriched LFP activity with low and stable EMG tone or Acc signal amplitude.

Experienced scorers first completed a training phase using 3 hours of simultaneously recorded LFP/EMG and LFP/Acc data compared against a consensus reference set to ensure proficiency in EMG and Acc scoring. This reference set was created by three experienced scorers who resolved all disagreements through discussion. Scorers were required to meet the following reliability criteria calculated versus the reference set before advancing to the experimental dataset: overall vigilance state accuracy > 90%, Cohen’s kappa > 0.80, and epoch-by-epoch agreement > 90%^11-20^. After meeting training criteria, scorers each scored the full experimental dataset. The full experimental dataset encompassed a total of 40 hours and 14 minutes which would be scored twice, once using LFP/EMG and once using LFP/Acc. Individual traces lasted between 47 and 82 minutes, with an average length of 65 minutes and 13 seconds. Traces were presented in a randomized order for every scorer; however, presentation did follow a sequential order of alternating EMG and Acc traces. Traces from the same mouse were never presented consecutively.

### Signal Processing and Analysis

All signal processing and analysis was performed in MATLAB 2025a (MathWorks) and OriginPro 2025b (OriginLab). LFP signals were downsampled to 2500 Hz. EMG signals were bandpass filtered at 300–1000 Hz and Acc signals at 0.5–100 Hz. The EMG filtering bandpass was chosen based on traditional field usage^5,9-10^. The bandpass for the Acc signal was set to encompass the frequencies of frequently encountered rodent behavior^30^.

For each individual epoch, signals were first bandpass filtered. The root means square (RMS) of the filtered signal was then computed for each epoch within its respective frequency band. To account for inter-recording variability in baseline signal amplitude and noise levels, RMS values were z-scored independently within each recording using the mean and standard deviation of all RMS values from that recording. A two-dimensional density plot of EMG (x-axis) vs Acc (y-axis) RMS z-scores were constructed by binning all pooled epochs into a 150 × 150 grid spanning −2 to 5 z-score units on both axes. Bin counts were normalised to the relative frequency of total epochs falling within each bin, allowing direct comparison of the distribution density across the two-dimensional space. Linear regression (x = EMG, y = Acc) was fit to the pooled epoch-level data using a least-squares polynomial fit (MATLAB polyfit), and Pearson’s r, R^2^, and the associated p-value were computed from the same pooled data. Density plot and regression procedure was applied separately to the subset of epochs on which EMG and Acc-based scoring agreed and to the subset on which they disagreed. Epochs were pooled across scorers, with each epoch labeled according to scorer and agreement status. Density plots arranged epochs by both scorer identity and agreement status to assess consistency of the EMG-Acc relationship across raters.

Epoch-by-epoch overall accuracy was computed as the percentage of epochs scored an identical state label by both EMG and Acc-based scoring out of all epochs in a recording. Cohen’s kappa was computed per state for each scorer-recording pair by measuring scorer agreement on a binary classification and then applying the standard kappa formula, κ = (P^o^ − P^e^) / (1 − P^e^), where P^o^ is actual agreement and P^e^ is the agreement expected bychance based on how frequently each scorer used each label. Kappa interpretation followed the established benchmark categories^31^ and described values from automated scoring programs (κ < 0 = poor, 0.01–0.20 = slight, 0.21–0.40 = fair, 0.41–0.60 = moderate, 0.61–0.80 = substantial, 0.81–1.00 = almost perfect agreement)^12,16-17,19-20^. In addition to overall (all-state) metrics, performance was also separately quantified for each vigilance state (Wake, NREM sleep, REM sleep) using F1 score, with Acc-based scoring treated as the prediction and EMG-based scoring treated as the reference. In F1 score, precision was defined as the proportion of epochs classified as a state by the Acc also classified as the same state by EMG-based scoring (TP/(TP+FP)). Recall was defined as the proportion of epochs classified as a state by EMG-based scoring that were also classified as a state by the Acc-based method (TP/(TP+FN)), where TP, FP, and FN denote true positive, false positive, and false negative epoch counts, respectively. F1 score was then calculated as the harmonic mean of precision and recall for each state: (F1 = 2 × (precision × recall) / (precision + recall))^32^.

Per-state comparisons required a strategy to appropriately account for potential differences between animals while maintaining scorer-level resolution in the final comparisons. The analytical hierarchy for per-state analyses proceeded as follows: First, for each individual scorer, an animal-level data set was created, comprising all epochs recorded from that mouse across all its recording sessions. Second, total accuracy, Cohen’s kappa, and F1 score were independently calculated for each animal’s pooled epoch set. Third, we calculated the mean of these per-animal metric values. This approach ensured that each animal contributes equally to each scorer’s final performance estimate, regardless of whether some animals provided longer or more numerous recording sessions than others. Consequently, the per-state bar graphs represent each scorer’s typical per-state performance while controlling for inter-animal variability.

To visualize the relationship between EMG and Acc z-scores and identify signal characteristics associated with scoring agreement, a two-dimensional scatterplot of EMG (x-axis) vs Acc (y-axis) RMS z-scores were constructed by binning all pooled epochs into a 150×150 grid spanning −2 to 5 z-score units on both axes. For computational efficiency and visualization clarity, scatterplot rendering was limited to a maximum of 3,000 randomly sampled points per heatmap using speed-mode in OriginPro 2025b, however, all statistical analyses were performed on complete unsampled datasets.

To quantify the degree of EMG-Acc signal divergence between “Agreed” and “Disagreed” epochs, we calculated delta R^2^ (ΔR^2^) as the difference in coefficient of determination between disagreed and agreed subsets (ΔR^2^ = R^2^ Disagreed – R^2^ Agreed) for each vigilance state. This metric captures the magnitude of change in EMG-Acc coupling strength as a function of scoring agreement status. For each vigilance state and the overall pooled dataset, we performed paired t-tests comparing R^2^ values between “Agreed” and “Disagreed” epochs across individual scorers (n = 7).

Data was considered statistically significant if p value was lower than α = 0.05, except for multiple comparisons where P values were compared with Holm’s adjusted α levels. Adjusted α are reported explicitly in the text. All data in text and figures are reported as mean ± standard error.

## RESULTS

### Characterizing signal agreement between LFP/Acc and LFP/EMG

A representative recording illustrates the raw LFP, EMG and Acc signals across the full extent of a sleep recording (Figure 1A). Zoomed insets highlight the characteristic LFP waveform of Wake, NREM sleep, and REM sleep states, confirming the correspondence between vigilance states (derived from EMG/LFP scoring) and physiologically distinct hippocampal activity patterns. Epoch-by-epoch RMS z-score traces for EMG (orange) and Acc (blue) co-varied closely throughout. Example scoring disagreements for one scorer concentrate in isolated epochs identified by red bars at the base of the figure.

**Figure 1.**
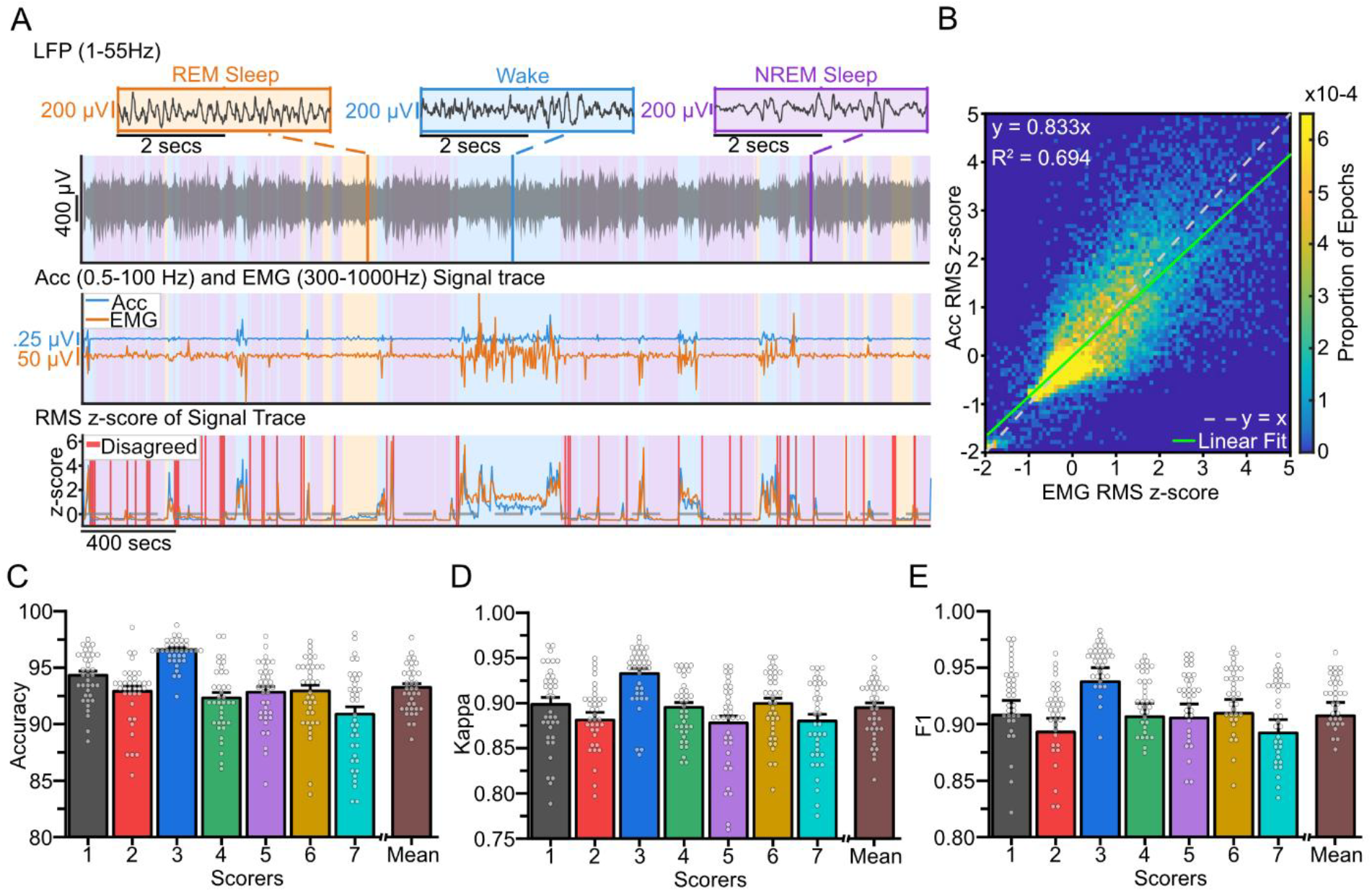
Signal comparison and scoring agreement. (A, Top) A representative single recording trace of raw LFP (1-55Hz) with vigilance state background shading (light blue = Wake, purple = NREM sleep, light orange = REM sleep) derived from EMG scoring and with insets showing characteristic single-epoch vigilance states. (A, Middle) A corresponding representative recording with a bandpass filtered EMG (300-1000Hz, orange) and Acc (0.5-100Hz, blue) signal trace. (A, Bottom) Corresponding epoch-by-epoch root-mean-squared (RMS) z-score signal traces for EMG and Acc along with red bars indicating epochs for which EMG and Acc scoring disagreed. (B) Two -dimension density heatmap of Acc (x-axis) vs EMG (y-axis) RMS z-scores per epoch of the filtered signal tace (colored per proportion of total epochs) with a green regression line (y = 0.833x + 0.000, R^2^ = 0.694, r = 0.833) and a y = x dotted line. (C) Mean overall accuracy (%) per scorer with individual scorer-recording trace points and mean scorer accuracy of 93.26. (D) Mean kappa score per scorer with individual scorer-recording data points and overall mean kappa score of 0.895. (E) Mean F1 score per scorer with individual scorer-recording data points together with mean F1 score of 0.91. Individual points represent single recordings.

Before determining the degree of agreement between Acc and EMG-based sleep scoring, we first sought to determine whether the two modalities co-varied consistently. To directly compare the EMG and Acc recording, we computed the z-scored RMS for both signals during each 4-second epoch and examined their relationship (n = 36,197 epochs). A two-dimensional density heatmap of EMG (x-axis) vs Acc (y-axis) z-scores revealed largely overlapping distributions for the two signals. A majority of datapoints were concentrated near the origin, indicating largely similar activity profiles of EMG and Acc across the recording (Figure 1B). This density plot also revealed a strong positive correlation (R^2^ = 0.694), confirming that Acc and EMG signals co-vary consistently.

Across all seven scorers, LFP/Acc based scoring achieved consistent and substantial epoch-by-epoch agreement with LFP/EMG (Figure 1C-E). Mean overall accuracy exceeded 90% for all scorers (Figure 1C), kappa was consistently above 0.80 (Figure 1D), and F1 scores exceeded 0.9 across all scorers and recording traces (Figure 1E), confirming low inter-scorer variability (mean accuracy across scorers = 93.26±0.33%; mean kappa = 0.895±0.0053; mean F1 = 0.91±0.012) and showing that Acc achieves substantial epoch-by-epoch agreement with EMG across all scorers.

### Agreement varies across vigilance states

Because the importance of EMG signal for sleep scoring varies across vigilance states, we hypothesized that discrepancies between EMG and Acc would also vary across states. To explore these potential differences, we further analyzed agreement separately for each state (Figure 2A-C). Total accuracy, mean kappa, and mean F1 scores were remarkably consistent across all three vigilance states, mirroring the overall pattern shown in Figure 1C-E. Accuracy exceeded 93% for both Wake and NREM sleep, with substantially higher performance for REM sleep classification. A one-way ANOVA showed significant differences in accuracy across sleep states (F(2,18) = 24.80, p < 0.0001). Post-hoc Holm-Bonferroni tests demonstrated that REM sleep scoring (98.47±0.23%) was significantly more accurate than both Wake (94.80±0.58%, p < 0.0001, vs Holm’s corrected α = 0.017) and NREM sleep (93.69±0.61%, p < 0.0001, vs Holm’s corrected α = 0.025), with no significant difference between Wake and NREM sleep accuracy (p = 0.13, vs Holm’s corrected α = 0.05, Figure 2A). Mean kappa values indicated strong inter-rater agreement for all states. A one-way ANOVA showed significant differences in kappa between vigilance states (F(2,18) = 6.87, p = 0.0061) and post-hoc Holm-Bonferroni tests indicated that REM sleep had significantly higher kappa (0.922±0.011) than Wake (0.879±0.0099, p = 0.011, vs Holm’s corrected α = 0.025) and NREM sleep (0.869±0.012, p = 0.0026, vs Holm’s corrected α = 0.017), with no significant difference between Wake and NREM sleep kappa (p = 0.52, Figure 2B). F1 scores followed a different pattern, remaining uniformly high across states and indicating strong classifier performance across all behavioral states. A one-way ANOVA showed no significant difference between vigilance states for F1 score (F(2,18) = 1.56, p = 0.24), indicating comparable classification performance regardless of vigilance state (Wake: 0.920±0.0052, NREM sleep: 0.940±0.008, REM sleep: 0.931±0.0093, Figure 2C).

**Figure 2.**
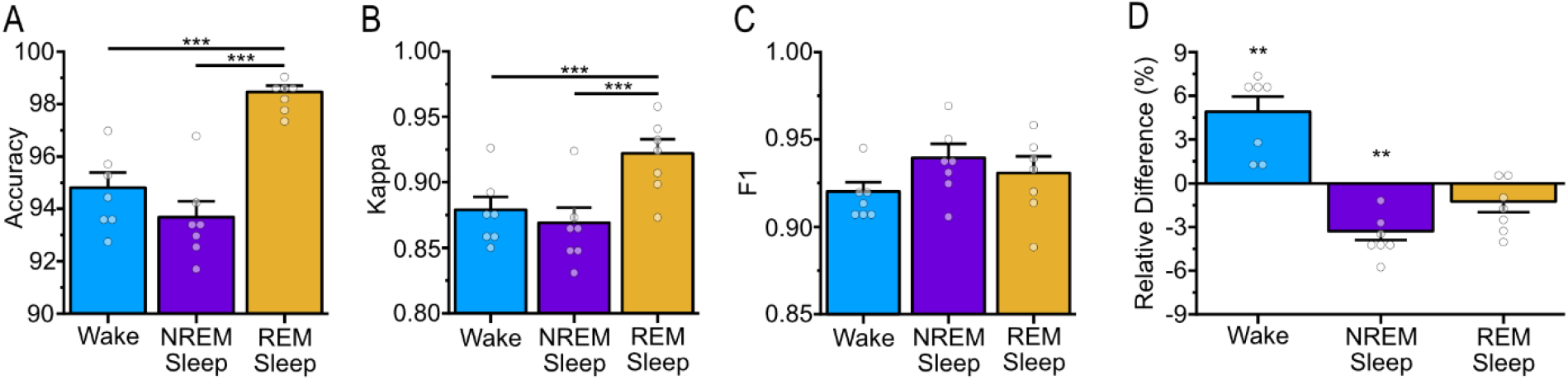
Per-state scoring agreement between EEG/EMG and EEG/Acc. (A) Mean accuracy (%) per-scorer during Wake, NREM sleep, and REM sleep. (B) Mean kappa score per-scorer during Wake, NREM sleep, and REM sleep. (C) Mean F1 score per-scorer during Wake, NREM sleep, and REM sleep. (D) Delta (EMG versus Acc) plot showing relative difference (%) of epoch counts for Wake, NREM sleep, and REM sleep. Individual data points represent each scorer.

Given that the agreement between the two scoring modalities was different between vigilance states, we next evaluated whether Acc-based scoring introduced systemic biases towards a particular vigilance state. We calculated the relative difference of vigilance state labeling between EMG and Acc, which showed a significant under-classification of Wake by the Acc (Figure 2D, - 4.92±1.03%, p = 0.0031) and significant overclassification of NREM sleep by the Acc (+3.29±0.61%, p = 0.0017), with number of epochs scored as REM sleep being similar between scoring methods (+1.29±0.73%, p = 0.14).

### Signal divergence between EMG and Acc predicts disagreement

To further understand whether disagreement between scoring from the two modalities stemmed from genuine difference in the movement-related signals themselves, we looked at the correlation between the RMS z-scores of each modality, similar to Figure 1B, but separated by agreement status and vigilance state (Figure 3).

**Figure 3.**
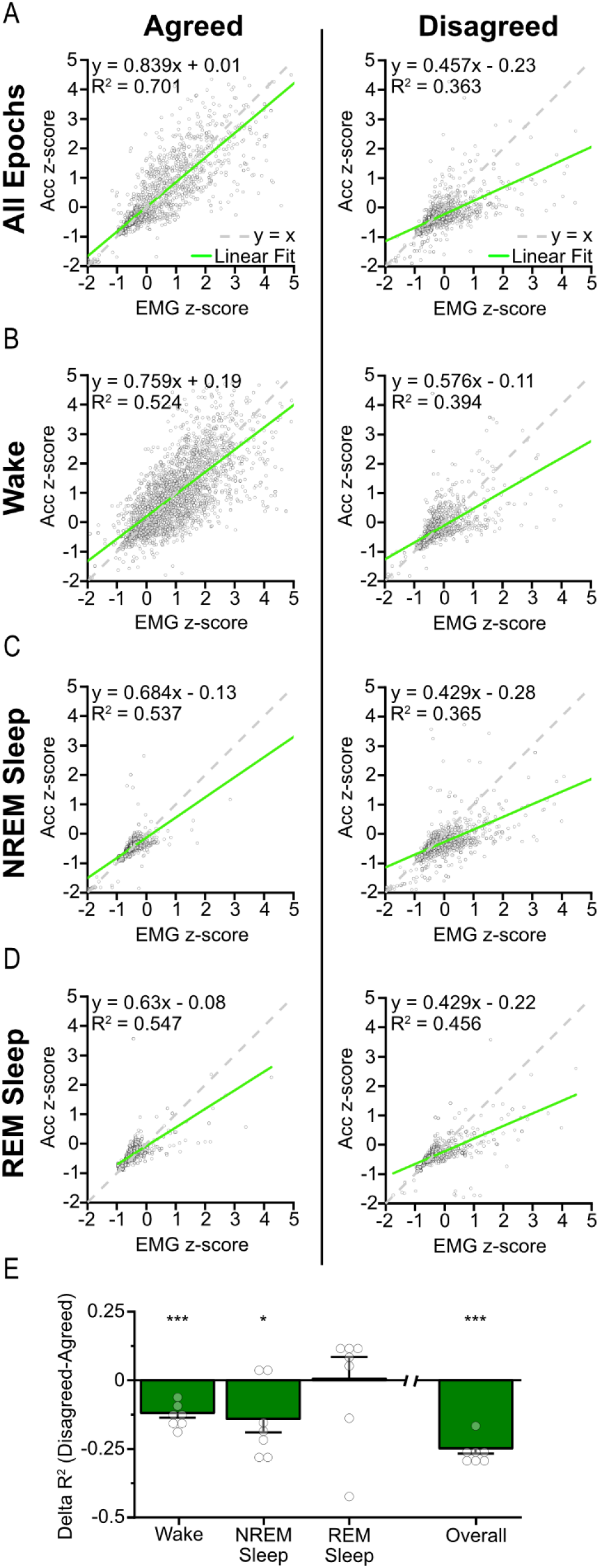
Signal divergence between EMG and Acc predicts scoring disagreement. (A-D) Density scatterplots comparing EMG RMS z-score (x-axis) to ACC RMS z-score (y-axis) for agreed (left column) and disagreed (right column) epochs across four sleep/vigilance states: (A) All Epochs, (B) Wake, (C) NREM Sleep, and (D) REM Sleep. Each example scatterplot displays a maximum of 3,000 randomly sampled epochs per EMG-ACC combination. Green regression lines indicate the linear relationship between EMG and ACC, with regression equation, R^2^, and sample size (n) displayed. Dashed gray lines represent unity (y=x) for reference. (E) Bar graph showing the delta R^2^ (R^2^ disagreed – R^2^ agreed) for each sleep state and overall pooled data. Individual data points represent each scorer.

We first considered all states together. Across a pool of all scorers and recording traces, 93.3% of epochs (n = 236,471) received identical scoring from both EMG and Acc, while 6.7% (n = 16,908) received different labels. For “Agreed” epochs, signal correlation was strong (Figure 3A, R^2^ = 0.701, p < 0.0001), with a tight density cluster along the diagonal. For “Disagreed” epochs (Figure 3A), the correlation was weaker (R^2^ = 0.363, p < 0.0001), with the density distributed more broadly and off diagonal comparative to “Agreed” epochs. We computed the difference in R^2^ between “Disagreed” and “Agreed” epochs for all scorers and found that R^2^ was significantly lower for “Disagreed” epochs (ΔR^2^ = -0.33±0.027, p < 0.0001, Figure 3E).

We further broke down the signal correlation between individual vigilance states (Figure 3B-D). For both Wake and NREM sleep, “Agreed” epochs again demonstrated notably higher R^2^ values (Wake: R^2^ = 0.524, NREM sleep: R^2^ = 0.537) compared to “Disagreed” epochs (Wake: R^2^ = 0.394, NREM sleep: R^2^ = 0.365, Figure 3B-C). In contrast, for REM sleep “Agreed” and “Disagreed” displayed relatively similar R^2^ values (Agreed: R^2^ = 0.547, Disagreed: R^2^ = 0.456, Figure 3D). To quantify the change in R^2^ values between “Agreed” and “Disagreed” epochs for each vigilance state (Figure 3E), we calculated delta R^2^ (R^2^ Disagreed – R^2^ Agreed). A one sample t-test revealed that R^2^ was significantly larger in “Agreed” epochs for both Wake (ΔR^2^ = -0.12±0.017, p < 0.0001) and NREM sleep (ΔR^2^ = -0.14±0.049, p = 0.029), while during REM sleep, the R^2^ for EMG-Acc correlation was similar between “Agreed” or “Disagreed” epochs (ΔR^2^ = +0.00±0.0795, p = 0.95). This indicates that as the signals from the two modalities diverge (low R^2^), the resulting scoring is more likely to disagree.

## DISCUSSION

This study provides the first direct comparison of LFP/EMG and LFP/Acc for manual sleep scoring in mice. Our results demonstrate that LFP/Acc-based scoring achieves substantial agreement with LFP/EMG, supporting the use of headstage accelerometry as a valid substitute to EMG recording. This conclusion relies on the consistently high accuracy, kappa, and F1 scores observed across all seven scorers at the overall epoch-by-epoch level.

The overall agreement metrics between EMG and Acc-based sleep scoring we report (accuracy > 90%, kappa > 0.8, F1 > 0.9) encourage Acc’s validity as these values are broadly consistent with benchmarks reported for automated and semi-automated LFP/EMG sleep-scoring tools in rodents. Typical reported human-versus-algorithm overall accuracy ranges from 80.24% to 97%^11-15,18-20^, and Cohen’s kappa values of approximately 0.75–0.94^12,17-20^. Importantly, our Cohen’s kappa comparing EMG and Acc also reaches similar levels or even exceeds that of recently published human-versus-human inter-rater agreement^33^. This study reported Cohen’s kappa for Wake between 0.91-0.95, NREM sleep between 0.86-0.94, and REM sleep between 0.78-0.93, placing all our agreement metrics towards the upper end of these ranges. Accordingly, using Acc instead of EMG does not seem to introduce more variability than that already inherent to available scoring methods.

These results are also consistent with the strong correlation between EMG and Acc RMS z-scores across all pooled epochs, showing that the two modalities are tracking a largely similar underlying physiology. Furthermore, we show when breaking this signal correlation down into “Agreed” or “Disagreed”, there is a substantially weaker correlation between EMG and Acc z-scores in “Disagreed” epochs compared to “Agreed” epochs. This demonstrates that disagreements, at least partially, arise when the two modalities diverge in their estimate of movement/activity level at the epoch level. Our results hint at REM sleep identification relying heavily on LFP for intra-hippocampal recordings, while Wake and NREM sleep rely more on movement/muscle tone provided from either an EMG or Acc, explaining the significant difference between Wake and NREM sleep, but not REM sleep.

Overall, this study shows that Acc-based scoring performs consistently well compared to EMG, with a strong identification of all vigilance states. However, we did uncover a small, but still notable, bias in Acc towards the labeling of NREM at the expense of Wake. This reinforces the observed state-dependent nature of EMG-Acc signal divergence. Disagreements seemed concentrated at behavioral state transitions, as exemplified in Figure 1A, suggesting that this ambiguity may also reflect an inherent challenge in precisely defining state boundaries using movement-based signals. However, future work addressing state-transition classification should further expand on this notion. This has an important practical implication as disagreements between LFP/EMG and LFP/Acc scoring cannot be resolved through improved scorer training, consensus procedures, or algorithmic refinement applied to the scorer level.

Several limitations should be considered when interpreting these findings, as this study utilized LFP recordings rather than full cortical EEG, which captures a more spatially distributed area of neural activity. LFP signals generally only reflect local intracranial electrical activity and may not fully capture the broad cortical signatures used traditionally in sleep scoring, so these findings may not translate exactly to multi-channel EEG recordings. Secondly, electrode placement is a critical variable which affects both LFP and EMG signal quality. Our findings reflect a specific hippocampal LFP and trapezius EMG placement, so different electrode configurations could yield different relationships between EMG and Acc. Lastly, our heavily conclusions rely on manual scoring by trained scorers, and all comparisons are reported from inter-rater agreement with no comparison to intra-rater agreement. While we have shown LFP based EMG to be a valid equal to Acc within the same scorer, we do not know if it is as valid as just using another manual scorer.

LFP/Acc-based sleep scoring represents a practical and well-validated approach that substantially reduces surgical complexity and risk while maintaining strong overall agreement with traditional EMG-based methods. The modest limitation in Wake-NREM sleep discrimination should be weighed against the considerable reduction in procedural burden, making LFP/Acc an attractive alternative.

## RESOURCE AVAILABILITY

The data that support the findings of this study are available from the corresponding author upon reasonable request.

## LEAD CONTACT

Requests for further information and resources should be directed to and will be fulfilled by the lead contact, Bénédicte Amilhon.

## ACKNOWLEDGMENTS

We thank Véronique Pellerin, Denise Carrier, and the animal facility team for support in animal care.

## FUNDING INFORMATION

DG was supported by the Bourse de la relève of the Faculty of Medicine of Université de Montréal (UdeM), the recruitment scholarship of the CHU Sainte-Justine Foundation and the recruitment scholarship of the Department of neurosciences of UdeM. JFH was supported by a Canada doctoral graduate scholarship from the Natural Sciences and Engineering Research Council of Canada (NSERC) and by a complement scholarship from the Fonds de recherche du Québec (FRQ). EL was supported by the recruitment scholarship of the CHU Sainte-Justine Foundation. TL was supported by a scholarship from the Natural Sciences and Engineering Research Council of Canada, the Catherine Fradette prestige doctoral award in biological science and neuroscience of the Université de Montréal, and the first Serge Rossignol prestige doctoral award of the Université de Montréal. AGC was supported by the recruitment scholarship of the Department of neurosciences of UdeM and the Canada Graduate Scholarships – Master’s of the Canadian Institutes for Health Research (CIHR). VM supported by the Canada Research Chair in Sleep Molecular Physiology. B.A. was supported by a Research Scholar salary award (Chercheur-boursier Junior 1 and Junior 2 from FRQS #253671 and 329843).

This work was funded by NSERC Discovery grant RGPIN-2018-06765 and CIHR Project grant #173244.

## AUTHOR CONTRIBUTION

Conceptualization: DG, GD, and BA; Investigation: DG, JFH, GD, and BA; Data curation: DG, JFH, EL, NL, TL, GD, and VM; Methodology: DG, JFH, EL, NL, TL, GD, and VM; Software: DG and GD; Formal analysis: DG and GD; Visualization: DG and GD; Writing – original draft: DG; Writing – review and editing: DG, JFH, NL, TL, GD, BA, and VM; Supervision: BA; Funding acquisition: BA.

## DECLARATION OF INTERESTS

The authors declare that they have no conflicts of interest.

## Notes

### Competing Interest Statement

The authors have declared no competing interest.

